# The Process Pathway Model of bacterial growth

**DOI:** 10.1101/553982

**Authors:** Daniel Biro, Ximo Pechuan, Maryl Lambros, Aviv Bergman

## Abstract

The growth profile of microorganisms in an enclosed environment, such as a bioreactor or flask, is a well studied and characterized system. Despite a long history of examination, there are still many competing mathematical models used to describe an output of the microorganisms, namely the number of bacteria as a function of time. However, these descriptions are either purely phenomenological and give no intuition as to the biological mechanisms underlying the growth curves, or extremely complex and become computationally unfeasible at the population level. In this paper, we develop the Process Pathway Model by modifying a model of sequential processes, which was first used to model robustness in metabolic pathways, and demonstrate that the Process Pathway Model encapsulates many features and temperature dependence of bacterial growth. We verify the predictions of the model against growth data for multiple species of microorganisms, and confirm that the model generates accurate predictions on temperature dependence of bacterial growth. The model has five free parameters, and the simplifying assumptions used to build the model are built upon biologically realistic notions. The Process Pathway Model accurately models a microorganism’s growth profile at an intermediate level of complexity that is computationally feasible. This model can be used as both an conceptual model for thinking about systems of bacterial growth, as well as a computational model that operates at level of complexity that is amenable to large scale simulation. This balance in accuracy and intuitiveness was accomplished by using realistic biological assumptions to simplify the underlying biology, which may point the way forward for future models of this type.

## 1 Introduction

The growth profile of microorganisms in an enclosed environment, such as a bioreactor or flask, known as batch culture, is a commonly used and well studied and characterized system [8,20]. It has applications to many fields, including food science, microbiology, experimental evolution, and bioreactor engineering [4,5,15, 24]. However, despite a long history of examination, there are still many competing mathematical models used to describe the output of the system, namely the number of bacteria as a function of time. Furthermore, these descriptions are either purely phenomenological, which give no biological intuition into the mechanisms underlying the growth curves, or extremely complex, becoming computationally unfeasible at the population level. The most common empirical models are the Logistic, Gompertz, van Impe, and Baranyi-Roberts, which describe growth profiles as concentrations over time for a population [2, 7, 23]. At the higher end of complexity, the full cell computational model of Karr *et al.* [13] predicts division time as a function of fundamental metabolic processes for a single low complexity species, *Mycomplasma genitalium*. Here we build off of a model of sequential processes developed by Kacser and Burns, originally used to model enzymatic robustness in metabolic pathways [11, 12], in order to develop a model of intermediate complexity between an empirical model and full cell metabolic integration. By adding temperature dependence to the simple model of Kacser and Burns [11], our model, termed the *Process Pathway Model*, encapsulates many features and temperature dependence of bacterial growth. Our results demonstrate that a relatively simple mechanistic model can be used to accurately describe and predict the dynamics of a complex biological system while maintaining biological relevance, computational tractability, and broad applicability.

Previous models of bacterial growth have been developed to predict bacterial growth rates as functions of time and temperature [2, 3, 9, 21, 22, 25]. Additional features that models are designed to predict are lag time, which is the time the bacteria spend in a stationary state before growing; carrying capacity, which is the maximal concentration that the bacteria grow to; and maximal growth rate, all as functions of variables such as temperature and pH. In addition to closed-form equations, differential equation models have also been applied to the problem of bacterial growth rates. The most common of these models is the Baryani-Roberts model [2, 3]. The equations of the model are integrated to obtain growth curves, and the predictions are typically better than those of closed-form equations, though parameters such as lag time, maximal growth rate, and carrying capacity have to be explicitly added to the differential equation models (see Supplementary Data: Fig. S4 for full comparison of models). Refinements and additions to the Baranyi-Roberts and van Impe models have had some success is predicting population level phenomena by approximating underlying processes, but these models remain highly phenomenological with little ability to extract biological insight into key parameters, such as the lag time [1, 14, 17–19, 25]. Other empirical models such as the Ratkowsky model, asymptote model, and hyperbola model describe a single aspect of bacterial growth, such as growth rate, carrying capacity, or lag, respectively, as functions of temperature, and are adaptable to numerous species of bacteria [21, 22, 26–28]. However, each empirical model is designed to explain only a single aspect of bacterial growth, such as density as a function of time, carrying capacity, or lag time, but no model has successfully integrated all of these features into a greater framework.

To further the understanding of the underlying phenomenon of bacterial growth and predict many key features of growth in a way that is computationally feasible, we introduce the Process Pathway Model. While abstract in nature, the model is rooted in the fundamental processes of biology, without becoming overly complex. The model is derived from a model of enzymatic robustness first put forward by Kacser and Burns [11]. However, rather than individual enzymes, gene expression and other cellular and physiological processes are modelled as being the underlying phenomena in the model. The model predicts key features of growth, specifically the lag time, maximal growth rate, and carrying capacity, as functions of temperature with similar accuracy to existing models of bacterial growth, but with more biological meaning and computational tractability. Additionally, the model can predict growth under fluctuating temperatures. Thus the model is useful on two levels; firstly as a model of simplifying biological assumptions to distill the most important abstractions involved in bacterial growth. Secondly the Process Pathway Model can be used as an accurate model to make predictions of the quantitative parameters involved in bacterial growth

## 2 Material and Methods

### 2.1 Mathematical Model

The Process Pathway Model consists of a chain of N processes, each processing an input and generating an output. Each process can be thought of as a series of physiological and gene regulatory activities. For example, one process could model the up regulation of gene expression in a particular metabolic pathway as a result of exposure to a novel resource rich environment. The totality of these processes, each representing a different cellular activity, comprise the network of cellular metabolism.

Each process in the chain of *N* processes is governed independently by Michaelis-Menten kinetics, resulting in a flux, *ϕ_i_* between processes given by:

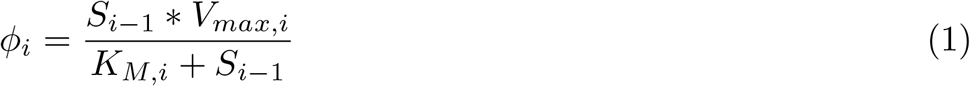

for *i* = 1, 2, 3, …, *N*, where *S_N_* is the concentration of the output of process *N*. For this model we assume no flux into *S*_0_ and the flux out of *S_N_* is given by a linear death rate. The dynamics for the variables *S_i_* is then given by:

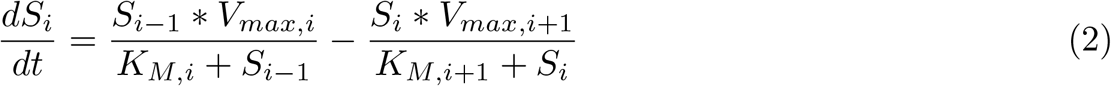

for *i* = 1, 2, 3, …, *N* − 1,

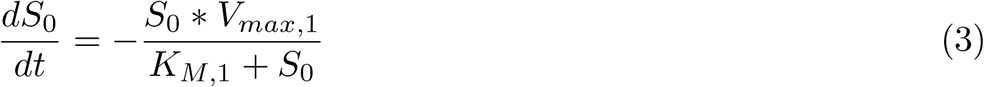

for *i* = and

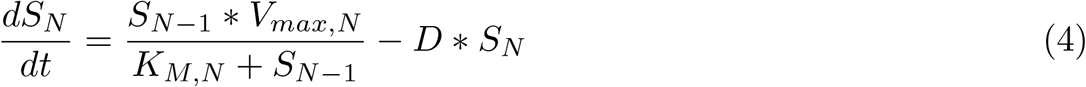

for *i* = *N* (Fig. 1 Panel A). *S*_0_ represents the concentration of the limiting resource, and is subsequently reduced as it is consumed by the process generating *S_1_*. The initial value of *S*_0_ is a free parameter, while the initial value of all other *S_i_* are set to 0 to model an initial state before growth has begun. The concentration of bacteria at time *t* is taken to be the value of the final process, *S_N_*(*t*), while the value of all other *S_i_*(*t*) represent the activity of the intermediate processes (Supplemental Figures: Fig. S1). Here, the final process is taken to represent the progress of the final metabolic pathway in the chain, in this case that of reproduction. The parameter *D* represents the natural death rate of the population. *V_max,i_* and *K_M,i_* here follow the same intuition as traditional Michaelis-Menten kinetics, where *V_max,i_* represents the maximal activity of each process, and *K_M,i_* is the concentration of substrate at which the process is at half of its maximal activity. In the original model of Kacser and Burns [11], *K_M,i_* were free to vary between enzymes; however here we take *K_M,i_* to be equal for all processes to reduce the number of free parameters without sacrificing significant accuracy. Other instatantiations of this model may benefit from the relaxation of this constraint.

**Figure 1.**
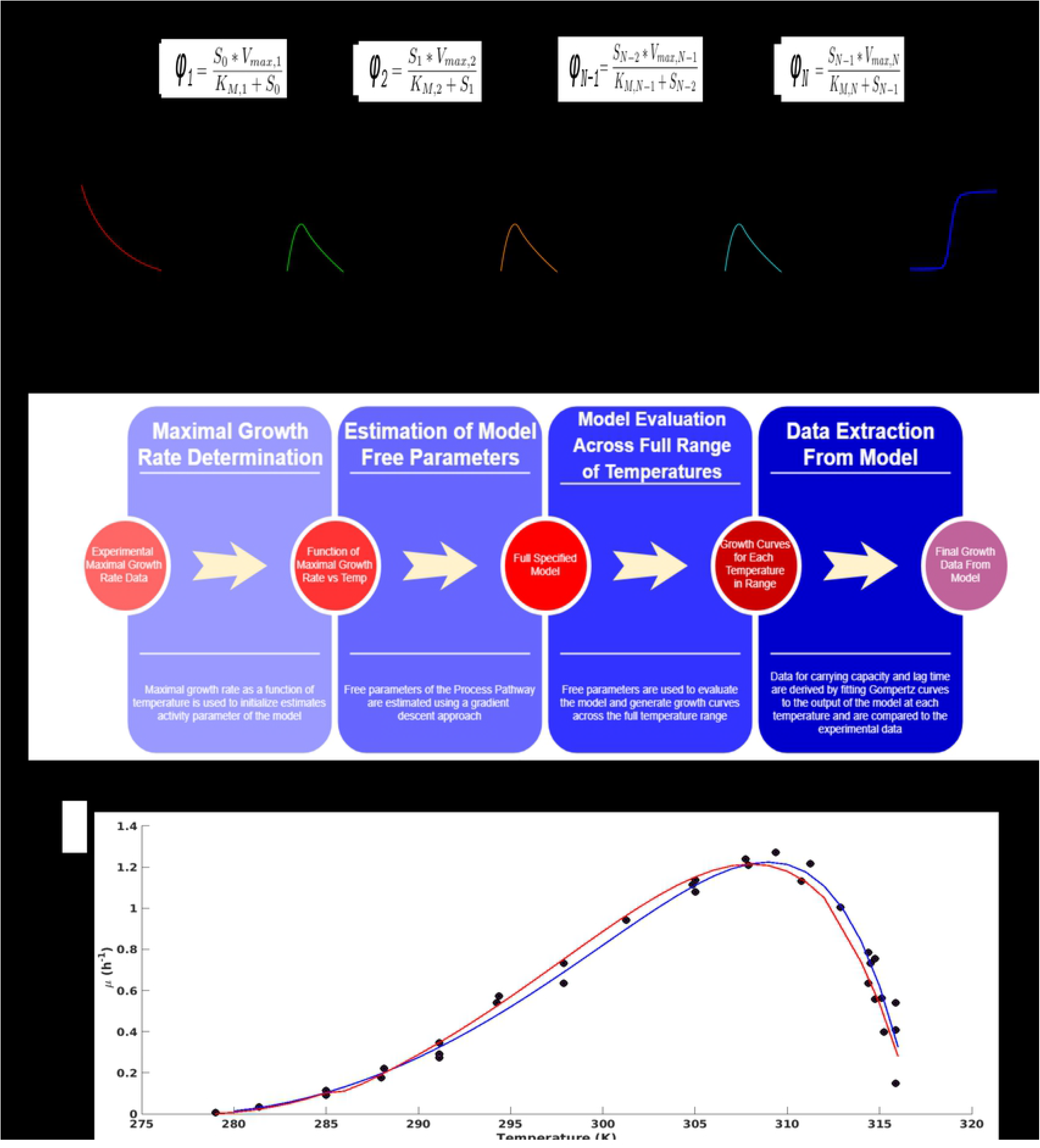

Temperature dependence of each process is incorporated by modelling a temperature dependence of the parameter *V_max,i_*. The functional form is a modified version of the function described by Daniels *et al*. [6], although the exact form of the temperature dependence did not significantly alter the results. The primary features of the temperature dependence that was salient for the model were the peak and the minima of the temperature dependence.

It is important to note here that while the equation for temperature dependence contain multiple parameters, they are not all free parameters, as the biological constraints effectively reduce the set of free parameters for temperature dependence of growth to a single value, visualized as the temperature of maximal growth. Thus, in all, this model has effectively three free parameters for defining bacterial growth; the maximal growth temperature defining the growth curve, *S*_0_, and *K_M,i_*. With the use of these parameters and the above assumptions which were used to construct the model, we can then generate predictions throughout the entire biokinetic range for the pertinent characteristics of bacterial growth.

In order to determine the optimal value for *N*, the number of processes in the chain, we compared experimental data from growth of *Escherichia coli* to the predictions of the model, yielding a optimal prediction of *N* = 8 (Supplemental Figures: Fig. S2). This is in line with predictions of the diameter of process networks in bacteria, as demonstrated by the whole cell computation model of Karr [13] utilizing 6 independent metabolic “compartments”. Furthermore, this result is robust for a wide range of values of *N* (see Supplemental Figures: Fig. S3), and these values are likely to be widely applicable, as network diameter scales with the logarithm of network size [10].

Applying the Pathway Process Model to data obtained from *E. coli* at a single temperature resulted in an accurate prediction of growth (Fig. 2 Panel A). The prediction was comparable to existing models of bacterial growth (Supplemental Figures: Fig. S4). Furthermore, the application of the Pathway Process model to data obtained of growth under a fluctuating temperature profile (Fig. 2 Panel B) fit the data as well as existing phenomenological models (Supplemental Figures: Fig. S5)

**Figure 2.**
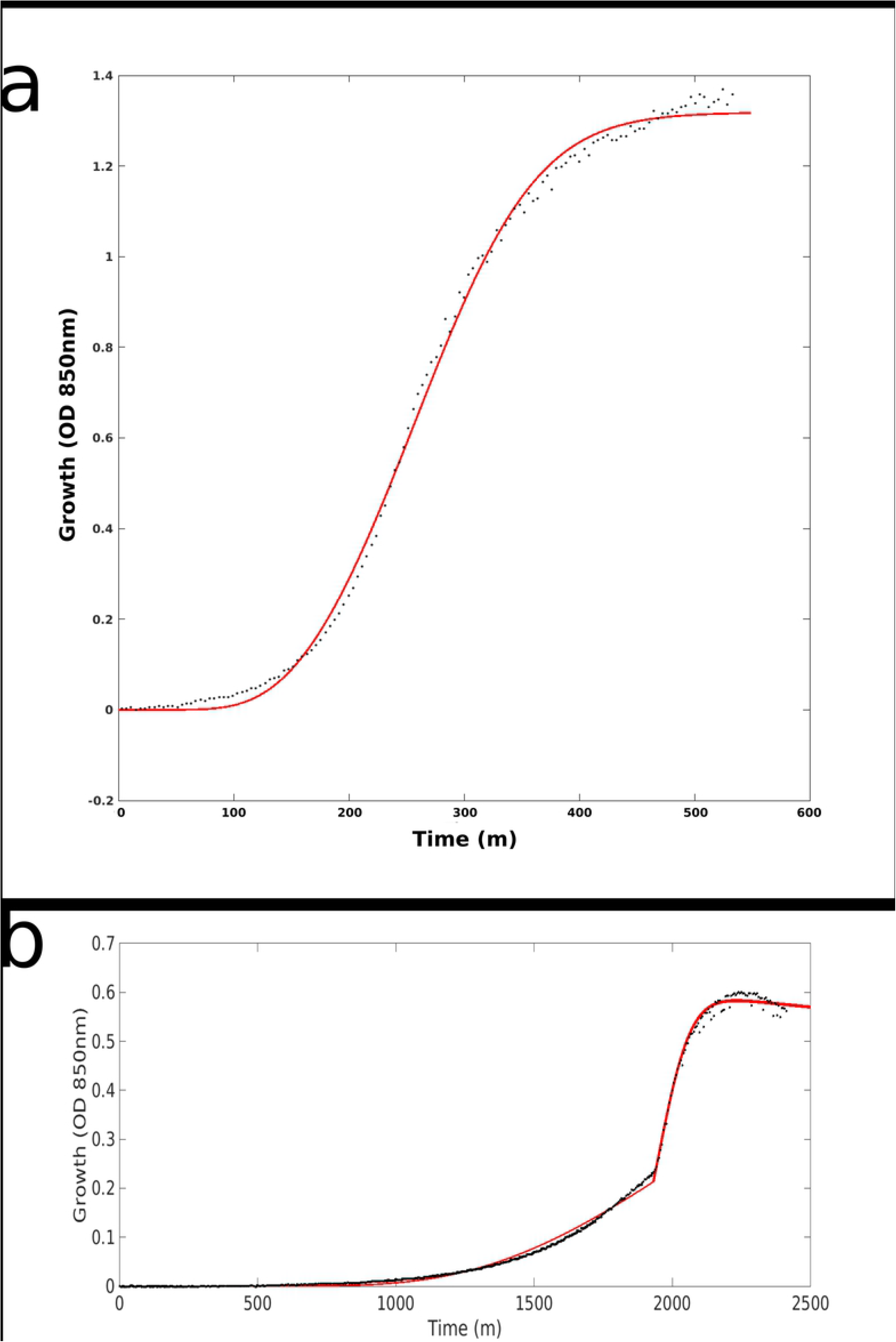

In order to test the ability of the model to predict the effects of temperature on the properties of bacterial growth, we determined a single set of parameters that best matched the data for growth rate, *μ*, for *Lactobacillus plantarum* data from Zwietering *et al*. [27] (Fig. 3 Panel A). The plot of the function for *V_max,i_*(*T*) using the parameters in Table S1 is shown in (Supplemental Figures: Fig. S5). It should be noted that the curve in this figure is derived experimentally, and is one of the few free parameters fed into the model. The results of the growth rate across the full dynamic range of the model are shown in Fig. 1 Panel C alongside the data raw data and the prediction from the empirical fit model. Zwietering *et al*.obtained values of *μ*, lag time, and carrying capacity by fitting bacterial count data to a Gompertz curve, so for consistency this is the same method used to extract parameters from the growth curves obtained from the Pathway Process Model. The algorithm used to estimate parameters and derive results is summarized in Fig. 1 Panel B.

**Figure 3.**
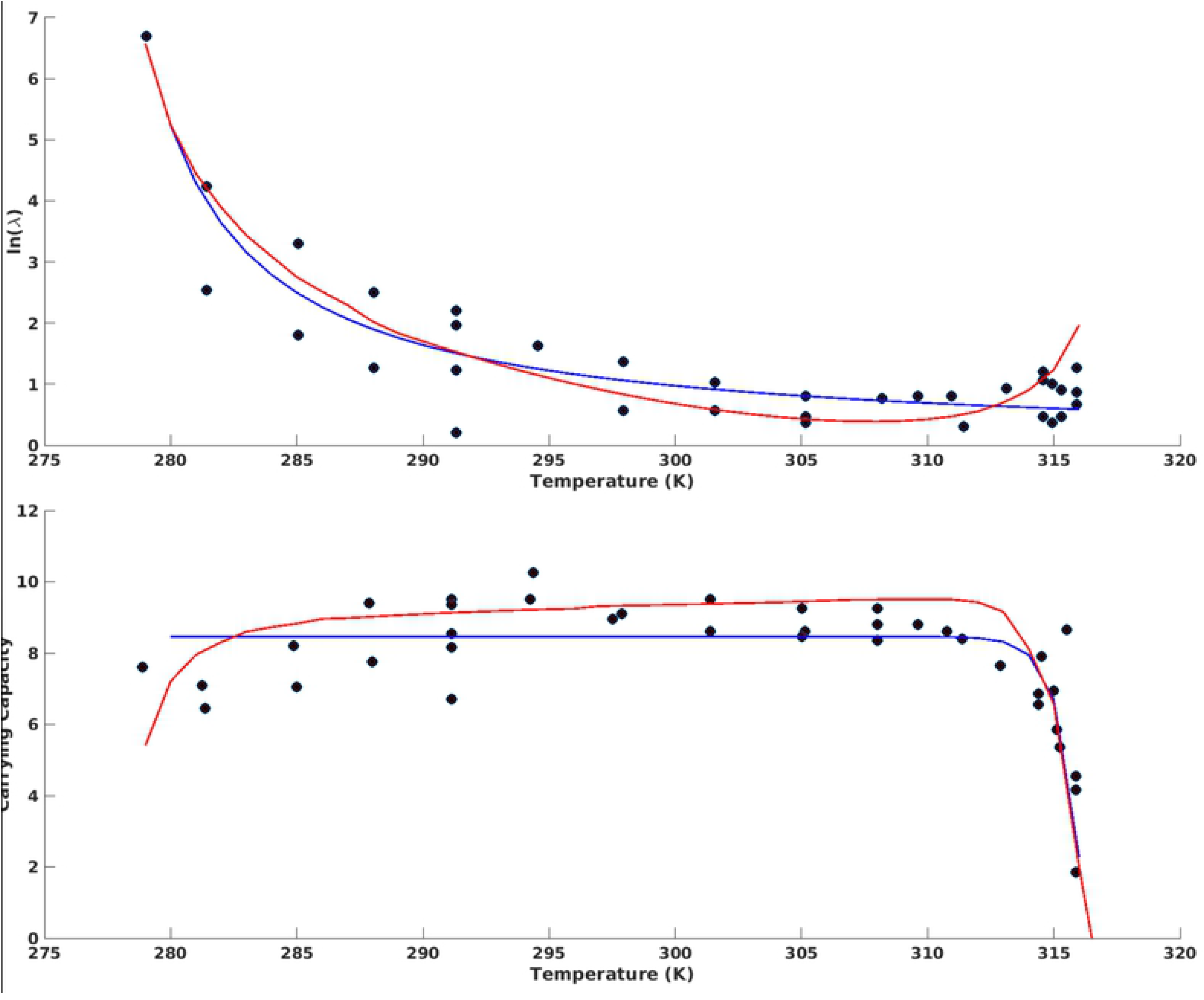

### 2.2 Computational Modelling

All computations were performed in MATLAB 2017a (Mathworks, Natick, MA). Solving of differential equations were performed using the function ode23s. Gompertz curves were fit using the Marquardt algorithm implemented in MATLAB.

Parameters for the temperature modelling were optimized by applying a gradient descent algorithm across all free parameters to minimise the least squares error between the predicted maximal growth rate, *μ*, and the data from Zwietering *et al*. Figure 2, except for *D*, which was chosen to be small. The least squares values were calculated by fitting a Gompertz curve to the output of the model integration, and extracting the parameter *μ* from this fit at each temperature. Once the parameters for temperature dependence were established using this procedure, the lag time and carrying capacity were similarly extracted from the Gompertz fit using the same parameters as for the maximal growth rate, *μ*. Parameters for all values are given in Table S1.

### 2.3 Data for Bacterial Growth Measurements

A derivative strain of *Escherichia coli* REL606 was used to seed the experiment. Bacteria were grown in M9 media with glucose supplemented to 4g/L. Tubes containing media were placed in BioSan LV Personal Bioreactor RTS-1C (Riga, Latvia), which controlled temperature to within 0.1°C. OD850nm measurements were taken by the RTS-1C at 2 minute intervals. To seed the culture, 100*μ*L of old culture was transferred to a new tube containing 20mL of fresh media.

Temperature switching and control was performed by the built in function of the Bioreactor RTS-1C. Temperature switching was programmed to occur after specified measurements of OD850nm occurred for the first time in a growth cycle. After temperature switching occurred, new temperature equilibrium was reached in approximately 30 minutes, which is less than the expected doubling time for the bacteria.

Each of the experiments started by inoculating the medium with 0.1 mL of an overnight culture grown at 37 ° in minimal media M9 with 10 % glucose. For the evaluation of the temperature dependence of the growth parameters, an additional day of acclimation was allowed [16].

### 2.4 Data for Growth Rate, Carrying Capacity, and Lag

Data for Figure 3 Panels A-C was used with permission from Figures 2, 5, & 6 of Zwietering *et al*. 1991. Data was originally collected from growth measurements of *Lactobacillus plantarum* in MRS media, with growth rates calculated by measuring CFU by titers, and parameters were calculated by fitting the data to a Gompertz curve.

## 3 Results

The two primary predictions of the model concerning the growth of bacteria over the entire dynamic temperature range, namely carrying capacity and lag time, are shown in Fig. 3 Panels A and B alongside the data from Zwietering *et al* [27]. In fact, aspects of the lag time and carrying capacity as functions of temperature are predicted by the model to a higher degree of precision than is seen in the empirical predictions. These results were obtained with minimal assumptions about the biology of the system and few free parameters.

The predictions for growth rate as a function of temperature generated by the process pathway model are equivalent to the empirical model in predicting the actual growth data (Fig. 3 Panel a). Similarly, the carrying capacity data derived from the process pathway model are shown to be similar to the empirical model in the middle of the growth range (Fig. 3 Panel C). Additionally, for carrying capacity, a slight decrease of the carrying capacity at low temperatures is predicted by the Process Pathway Model but not by the Ratkowsky asymptote model [27]. This results is shown both qualitatively and quantitatively in the data.

Additionally, for the lag time, the slight increase in lag time at higher temperatures was not accurately captured by the hyperbola model of lag time [27] but is predicted by the Process Pathway Model (Fig. 3 Panel B). Again, this result is shown both quantitatively and qualitatively as a prediction of the process pathway model and in the empirical data, but is not expected or shown in the traditional empirical models of bacterial growth.

The results here are derived from a model that takes in effectively five free parameters, and uses simplifying assumptions about the dynamics of bacterial growth in order to predict the biokinetic proprieties of growth throughout the viable temperature range.

## 4 Discussion

Bacterial growth is a highly complex phenomenon with many influences and complex behaviors. While there are numerous models describing growth, all are either empirical or detailed to the extent of being computationally burdensome, and none to our knowledge incorporate a framework based on simple abstractions of fundamental metabolic processes. Additionally, each empirical model is designed to explain only a single aspect of bacterial growth, such as density as a function of time, carrying capacity, or lag time, but no model has successfully integrated all of these phenomenon into a greater framework. Here we have put forward a model that encapsulates many of the pertinent features of bacterial growth, particularly in regards to temperature sensitivity, while being computationally tractable enough to be used for population level modelling. While biologically, the different phases of growth involve numerous different mechanisms and pathways, we were able to successfully abstract them away into their fundamental contributions in this model. This was surprisingly able to be successfully done with a single set of parameters for all phases. Many studies, including experimental evolution and protein structure and stability studies take interest in the effects of temperature on metabolic processes. Our model can successfully predict the above phenomenon with minimal assumptions and few free parameters. We reiterate that this model effectively has three free parameters, and the simplifying assumptions used to build the model are built upon biologically realistic assumptions.

One of the interesting and counter-intuitive insights gleaned from this model is the relationship between lag time and maximum growth rate. Previous models have treated these two variables as independent entities, though perhaps correlated, can be independently modelled. However a direct result of the process pathway model is that the lag time is simply a result of the rate of process activity, *V_max_* of each individual process. While this intuitively expected to be the case for the maximum growth rate, the more interesting result is that it is also the case for the lag time, which has been traditionally modelled as a phenomenon in its own right.

While this model is not a comprehensive model of bacterial metabolism or reproduction, we believe that it represents a form of a ‘minimal model”, where the pertinent features of the metabolic processes involved in reproduction are included, but the extraneous features are abstracted away. We have attempted to keep the number of free parameters to a minimum as to not overfit, while still being able to fit the data as accurately as models that do not have a basis in the fundamental biology. As such, this model represents an example what the authors feel is the appropriate abstraction of biological systems that renders them conducive to mathematical modelling and quantification while still retaining fundamental biological intuition.

